# The influence of canopy radiation parameter uncertainty on model projections of terrestrial carbon and energy cycling

**DOI:** 10.1101/618066

**Authors:** T. Viskari, A. Shiklomanov, M.C. Dietze, S.P. Serbin

## Abstract

Reducing uncertainties in Earth System Model predictions requires carefully evaluating core model processes. Here we examined how canopy radiative transfer model (RTM) parameter uncertainties, in combination with canopy structure, affect terrestrial carbon and energy projections in a demographic land-surface model, the Ecosystem Demography model (ED2). Our analyses focused on temperate deciduous forests and tested canopies of varying structural complexity. The results showed a strong sensitivity of tree productivity, albedo, and energy balance projections to RTM parameters. Impacts of radiative parameter uncertainty on stand-level canopy net primary productivity ranged from ~2 to > 20% and was most sensitive to canopy clumping and leaf reflectance/transmittance in the visible spectrum (~400 – 750 nm). ED2 canopy albedo varied by ~1 to ~10% and was most sensitive to near-infrared reflectance (~800 – 1200 nm). Bowen ratio, in turn, was most sensitive to wood optical properties parameterization; this was much larger than expected based on literature, suggesting model instabilities. In vertically and spatially complex canopies the model response to RTM parameterization may show an apparent reduced sensitivity when compared to simpler canopies, masking much larger changes occurring within the canopy. Our findings highlight both the importance of constraining canopy RTM parameters in models and valuating how the canopy structure responds to those parameter values. Finally, we advocate for more model evaluation, similar to this study, to highlight possible issues with model behavior or process representations, particularly models with demographic representations, and identify potential ways to inform and constrain model predictions.

**Key points:**

- Uncertainties in vegetation radiative parameter affect carbon, water and energy balances in ecosystem models.
- Radiative properties can lead to significant changes in demography and canopy structure even if aggregate model responses appear unchanged

## Introduction

Terrestrial vegetation regulates the global carbon cycle by taking up atmospheric CO_2_ through photosynthesis and releasing a smaller amount of CO_2_ back to the atmosphere through autotrophic and heterotrophic respiration (Field et al., 2007). In addition, seasonal and inter-annual vegetation dynamics drive a large fraction of the surface energy balance through regulating how solar energy is reflected back to the atmosphere (Fischlin et al., 2007). Importantly, variation and changes in surface albedo (through succession, disturbance, or land management) directly change the amount of radiation absorbed by the terrestrial biosphere, which has a first-order influence on global climate (Bonan, 2008). The interaction between available radiation, canopy architecture and optical properties drives a number of key processes including plant photosynthesis and surface water and energy cycling (Swann et al., 2010; Swann et al., 2012). These interactions are not only dependent on individual tree properties but also the canopy composition (Amiro et al., 2006). The radiation profile within the canopy is central to all of these processes. Understanding the complex relationships between radiation capture and utilization, with related processes such as carbon uptake, is necessary to inform and understand biosphere feedbacks to climate change.

Terrestrial Biosphere Models (TBMs), which combine the biophysics of a land surface model with representations for vegetation dynamics and biogeochemistry, are our primary tool for studying and projecting the impacts of current climate variability, disturbance, and on-going global climate change on the fluxes and storage of terrestrial carbon, energy, and water across space and time (Dietze and Latimer, 2011; Purves and Pacala, 2008; Fisher et al., 2014). We rely crucially on TBMs to test alternative hypothesis about ecosystem processes, project future ecosystem states, and understand how different processes and associated uncertainties affect those projections. There are numerous available TBMs (Schwalm et al. 2010; Dietze et al. 2011; Fisher et al. 2014), each representing different degrees of complexity, making different assumptions, and developed at different sites using different datasets for calibration, parameterization and validation. As a consequence, a number of key issues remain in the representation of surface dynamics, energy balance, and carbon fluxes and stocks, such as how best to represent different canopy layers within a model (Alton et al. 2007; Dietze and Moorcroft. 2011; Schaefer et al 2012; Keenan et al. 2012a; Friedlingstein et al. 2014). Furthermore, observations indicate that this forest complexity affects how the canopy responds to changes in vertical radiation profile (Hardiman et al., 2011, Stuart-Haentjens et al., 2015; Fahey et al., 2016), which is currently not captured by traditional models. Thus, as TBMs are developed towards more detailed descriptions of plant demography (Fisher et al. 2015; Fisher et al., 2017), it also becomes more important to understand the associated uncertainties and impacts of the radiation transfer schemes in order to better guide and evaluate this model development (Medvigy et al., 2009; Fisher et al., 2015; Weng et al., 2015).

To date, however, there has been relatively little focus on the parameterization and uncertainties surrounding radiative transfer in TBMs. The biophysical parameters required for properly simulating canopy photosynthesis, phenology, water conductance and surface energy balance (Sellers et al. 1997; Field et al. 2007; Bonan et al. 2011; Bernacchi et al. 2013; Rogers et al., 2017), are often poorly constrained in TBMs (e.g. Zaehle et al. 2005; Alton et al. 2007). In addition, the associated uncertainties are often ignored when examining model projection uncertainties and identifying model development needs (e.g. Alton et al., 2007; Keenan et al, 2012b). As the available radiation for photosynthesis is a driving force in canopy dynamics and competition, it is expected that allowing the canopy properties to vary would result in increased variability in simulated carbon, water, and energy fluxes (e.g. Alton and North, 2007). However, there could also be a number of compensating biophysical feedbacks. For example, altering light penetration should lead to changes in leaf and soil temperatures, which could change transpiration and evaporation, respectively, as well as total ecosystem respiration rates (Atkin and Tjoelker, 2003) and decomposition (Moore, 1986) thus indirectly affecting water, carbon cycling, and nutrient availability. Additionally, radiation is a limited resource within the canopy and a driver of competition and succession, particularly in dynamic vegetation models (Fisher et al., 2017), which suggests that the canopy response to changing radiative parameters might be more complex than a sum of individual parts (Dietze and Clark, 2008). Thus model development efforts should also consider these important and interconnected dynamics and examine multiple canopy outputs at varying scales as simply focusing on total canopy outputs can be misleading.

The objectives of our study are as follows: 1) To determine the impact of canopy radiative transfer (RT) parameter variation on the projection of C, water and energy fluxes between forests and atmosphere and 2) characterize how canopy structure affects RT uncertainty. These objectives allow us to evaluate which properties related to canopy radiative transfer need to be considered during model development activities, particularly those focused on improving the representation of the canopy radiation regime and associated processes representing carbon, water, and energy fluxes and stores. As the focus here is not on model validation, but rather on the internal model dynamics, the results will not be compared to observed time series. Our analysis was done with the Ecosystem Demography (ED2) model (Medvigy et al., 2009), a physiologically based forest canopy model that contains sophisticated cohort scaling of vegetation biomass together with a multi-layer land-surface scheme (Walko et al., 2000; Medvigy et al., 2009) and which contains a version of the two-stream radiative transfer model (Sellers, 1985) common to TBMs. Focusing on ED2 allows us to examine how uncertainties in canopy RT within a demographic TBM impacts model projections relevant to other comparable models (e.g. LPJ-GUESS; Smith et al., 2001; CLM(ED); Fisher et al., 2015). While we focus on a single model in this study, we also provide the framework for similar examinations with other models to assess how vulnerable they are to radiative parameter uncertainties.

## Methods

### 2.1 Model Description

For this study we utilized the Ecosystem Demography model v2.2 (ED2), a mechanistic TBM that contains a sophisticated approach for the scaling of forest dynamics from individuals to landscapes (Moorcroft et al., 2001; Medvigy et al. 2009; Medvigy and Moorcroft 2012). Within ED2, trees of similar size and properties within a patch are grouped together and represented by idealized cohorts in order to keep computational costs realistic. A sophisticated stage- and age-based approximation is used to scale up plant and stand level dynamics to regional-scale projections. ED2 also represents biophysical components common to land surface models (LSMs), which solve the surface energy and moisture budget, while also containing plant ecophysiology and CENTURY-style (Parton, 1996) soil biogeochemistry. As a result, ED2 can explicitly resolve processes across a range of temporal scales that span from instantaneous fluxes of heat, moisture, and energy to the centennial-scale dynamics of succession and vegetation migration. For this work we used the development version of ED2 (https://github.com/EDmodel/ED2, commit a4445f6)

Within ED2, the canopy RTM describes the penetration of incident (direct and diffuse) radiation through the canopy, which in turn drives photosynthesis, surface energy balance, and competition between vegetation cohorts. Although ED2 does not represent canopy horizontal heterogeneity in radiative properties (i.e. 1D RTM), it does represent multiple cohorts of trees of different sizes, stem densities, and plant functional types within a single patch. Cohorts of trees in ED2 have a height, leaf area, and canopy radius determined allometrically from the diameter at breast height (DBH), using the internal allometry within ED2 for each PFT (Dietze et al., 2008). Radiation in the canopy is represented in a multiple scattering model that is solved in a matrix solver, with leaf angle, transmission, reflectance, and emissivity set on a PFT basis. Radiation calculations also depend on topography, defined in ED2 as the slope and aspect of the site. In total there are three canopy solvers: direct shortwave (SW), diffuse SW, and longwave (LW) radiation. By default the soil optics in ED2 are fixed to a specific PAR and NIR range albedo, and in this study we used dry soil albedos of 0.25 and 0.39 for the PAR and NIR, respectively.

The sophisticated canopy scaling and cohort representation within ED2 enables the examination of how optical parameters interact with plant competition to affect the canopy and output uncertainties. As a result, this enabled us to examine both how the radiative properties affect the canopy at a single point of time, and how the canopy changes over time due to the change in the vertical radiation profile. Furthermore the basic RTM within ED2 (independent of canopy representation) uses the two-stream approach (Sellers, 1985) to model the interaction of light with the vegetation canopy, which is a common approach for TBMs. Thus even as we focus on ED2-RTM here, the results from these tests do also indicate how other TBMs using this RTM are affected by the uncertainties in radiative parameters.

### 2.2 Two-stream representation of canopy radiative transfer

Within the remote sensing and ecosystem modeling communities, the representation of canopy radiative transfer can vary considerably. Several prominent TBMs [e.g. JULES (Best et al., 2011; Clark et al., 2011), CLM (Lawrence et al., 2011; Lawrence and Fisher, 2013)] utilize the two-stream approach based on Sellers (1985). In this approach the fundamental calculations of the radiation regime depend on a few central parameters. The single scattering and backscattering coefficients for the leaf and wood within-canopy scattering elements describe the fraction of radiation scattered towards the surface and sky, respectively, and are calculated from the reflectance and transmittance spectra of the corresponding materials. These coefficients vary significantly across the shortwave infrared spectrum (i.e. 350 – 2500 nm), but for practical purposes are generally defined for two radiation wavelength ranges: Photosynthetically active radiation (PAR; radiation wavelengths of 400-800 nm) and infrared radiation (NIR; radiation wavelengths of 800-2500 nm). The wavelength bands used to calculate single scattering and surface albedo matters to the overall energy balance calculations since longer wavelengths reflect less light than the NIR and will lower the overall average albedo (Henderson-Sellers and Wilson, 1983). However, it is often difficult to identify the exact spectral range used to parameterize PAR and NIR reflectance and transmittance in models and the origin of the estimates of these values for each PFT.

In addition to the spectral properties of wood and leaf scattering elements, the canopy radiation profile also depends on the distribution of these elements in 3D space. A typical two-stream approximation of canopy radiative transfer assumes leaves are infinitesimally small, non-overlapping, and spread randomly through the canopy (i.e. “turbid medium”), and that leaf angles follow a spherical distribution. This allows the three dimensional radiative transfer problem to be approximated as a 1D (vertical) problem. To account for more realistic canopy geometry, two additional geometric parameters can be incorporated to the two stream RTM: canopy clumping factor and leaf orientation factor. The canopy clumping factor (Ω) describes the non-random distribution of foliar elements in a plant canopy (Nilson, 1971; Kucharik et al., 1997) and ranges from 0 (leaves tightly clumped together) to 1 (leaves homogeneously distributed with no clumping). Leaf orientation factor (*a_orient_*), in turn, establishes the average angle between the radiation and leaf surface (Josepoh et al., 1996) with values from −1 (all leaf surfaces are parallel to the direction of the radiation) to 1 (all leaf surfaces are perpendicular to the direction of the radiation).

The optical ‘thickness’ of each layer depends on the cohort leaf area index (LAI) and wood area index (WAI). In ED2, cohort LAI is a product of cohort stem density (*N_cohort_*), cohort foliar biomass (*B_Leaf,cohort_*) and specific leaf area (*SLA)*:

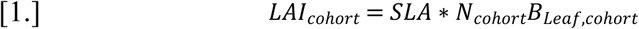

Cohort WAI is calculated according to Hormann et al. (2003):

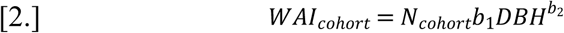

Where *DBH* is the trunk diameter at breast height and both *b_1_* and *b_2_* are PFT specific coefficients. With respect to radiation transfer, total area index (*TAI*) is a sum of effective LAI (*eLAI*), which is weighed by the clumping factor (*Ω*), and WAI.

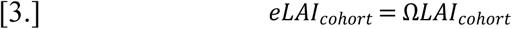

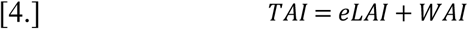

The location of a cohort layer in the RT scheme is determined by the cohort height. In ED2, cohorts interact with the radiation in layers proceeding from the tallest cohort to shortest cohort independent of how dense the actual vegetation within the canopy is. As each layer will absorb radiation, this reduces the amount of available radiation to the lower cohorts. Additionally, due to scattering within the canopy the upper layers can also receive radiation back from lower cohorts due to reflection. In ED2, the whole cohort is assumed to be in the same layer.

The cohort height is a function of the cohort DBH according to equation

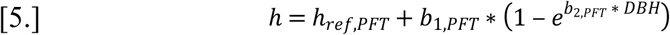

where *h_ref,PFT_* is an initial reference height for the given PFT and both *b_1,PFT_* and *b_2,PFT_* are PFT-specific height coefficients

The current ED2 RTM is a version of a classic two-stream RTM with each cohort representing a canopy layer based on their height order. In ED2, the single scattering (*a_scattering_*) and backscattering (*a_backscattering_*) coefficients are calculated as

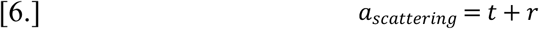

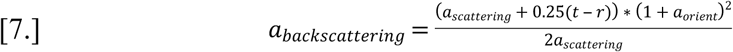

Where *t* is the transmission coefficient, *r* the reflective coefficient and *a_orient_* the orientation factor coefficient. The equation is the same, although with different coefficients, for both PAR and NIR ranges as well as for leaf and wood. Once all the area indices as well as scattering and backscattering coefficients have been determined, ED2 solves the fraction of radiation absorbed by each layer with linear equation solver. From these results, the total canopy albedo is then determined.

### 2.3 Experimental setup

All the analyses here were conducted with the Predictive Ecosystem Analyzer (PEcAn, LeBauer et al. 2013; Dietze et al. 2014) framework, which is an open-source scientific workflow and ecoinformatics toolbox that manages the flow of information in and out of TBMs and facilitates formal uncertainty quantification, data assimilation, and model benchmarking (https://pecanproject.github.io/).

The core component of the PEcAn framework used in this study was the uncertainty quantification (UQ) module, which allowed us to breakdown the impact of parameter uncertainty on model output (LeBauer et al., 2013; Dietze et al., 2014). In PEcAn the UQ information is provided as: 1) Coefficient of Variation (CV), which is the parameter standard deviation normalized with the mean parameter value. It is used to represents parameter uncertainty as it both generally indicates how large the uncertainty is and allows comparison between different parameter uncertainties affecting the system; 2) Elasticity, which is the change in the output variable in response to the change in parameter value. It is an indicator of output sensitivity to parameters and again allows comparison on how different parameters impact the canopy. It should be noted that the elasticity does not provide information on what the relation between the output and parameter is, though; 3) and Variance Decomposition (VD), which is the product of parameter uncertainty and variable sensitivity. This represents the total contribution of the parameter to model output uncertainty and allows isolating which parameters affect the current uncertainty the most. The reason this information isn’t sufficient by itself is that it does not differentiate if this impact is more due to the general model sensitivity to the parameter or because the initial parameter value itself is that uncertain. Additionally it is important to note these UQ only applies to the parameter impacts and does not represent the whole model uncertainty as this does not capture the structural uncertainties.

The impact of ten separate canopy RT parameters were studied here: Leaf PAR reflectance/transmittance (LR_PAR_ and LT_PAR_, respectively), leaf NIR reflectance/transmittance (LR_NIR_ and LT_NIR_, respectively), wood PAR reflectance/transmittance (WR_PAR_ and WT_PAR_, respectively), wood NIR reflectance/transmittance (WR_NIR_ and WT_NIR_, respectively), clumping factor and orientation factor. The parameters, the associated abbreviations and prior distributions are listed in Table 1. The PAR and NIR ranges for wood reflectance were derived from Asner (1998) and measurements of deciduous hardwood species in the Upper Midwest, US. These distributions are shown in Figure 1. For the leaf-level optical properties we derived the prior distributions from leaf spectra generated by the PROSPECT 5 model (Jacquemoud and Baret 1990, Feret et al. 2008) parameterized following the prior distributions on leaf biophysical parameters (number of mesophyll layers and chlorophyll, carotenoid, water, and dry matter contents) from Shiklomanov et al. (2016).

**Fig 1:**
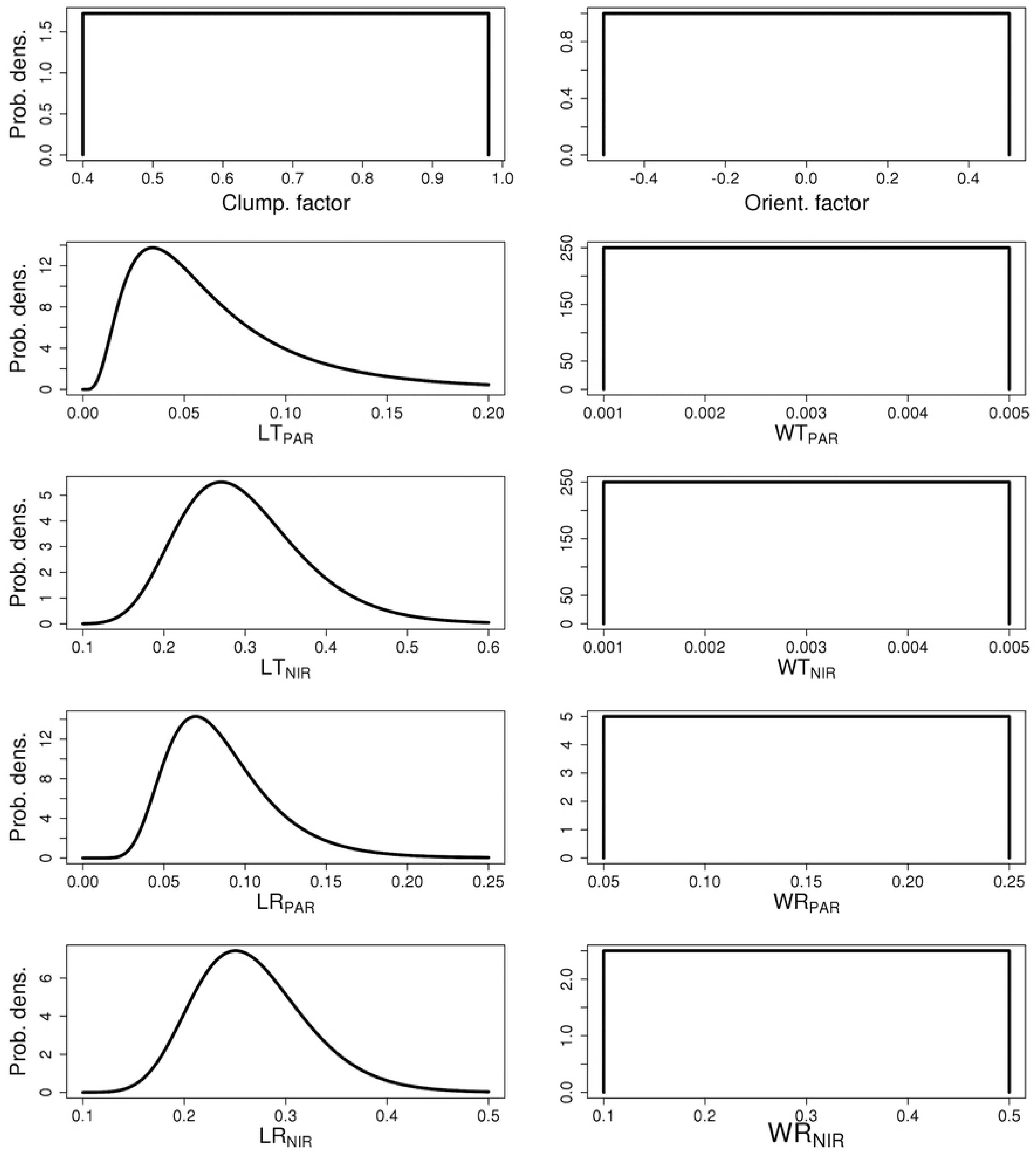
Prior probability distributions for chosen parameters

**Table 1:** Table of parameter used in these experiments

|  |  |  |
| --- | --- | --- |
| cf | Clumping factor | Unif(0.4,0.98) |
| Of | Orientation factor | Unif(-0.5, 0.5) |
| LT<sub>PAR</sub> | Leaf transmission in the PAR range | Lognormal(-2.9176, 0.6729) |
| LR<sub>PAR</sub> | Leaf reflectance in the PAR range | Lognormal(-2.5367, 0.3748) |
| LT<sub>NIR</sub> | Leaf transmission in the NIR range | Lognormal(-1.241, 0.2588) |
| LR<sub>NIR</sub> | Leaf reflectance in the NIR range | Lognormal(-1.3402, 0.2096) |
| WT<sub>PAR</sub> | Wood transmission in the PAR range | Unif(0.001, 0.005) |
| WR<sub>PAR</sub> | Wood reflectance in the PAR range | Unif(0.05, 0.25) |
| WT<sub>NIR</sub> | Wood transmission in the NIR range | Unif(0.001, 0.005) |
| WR<sub>NIR</sub> | Wood reflectance in the NIR range | Unif(0.1,0.5) |

The experiments were run with different combinations of Early-, Mid- and Late-successional Temperate Hardwood broadleaf tree PFTs. While the radiative parameter distributions in Fig. 1 were the same for all PFTs, the parameters guiding carbon allocation and photosynthetic processes differ between these PFTs. The PFT descriptions, including species, and parameter distributions are available through the BETY database (http://betydb.org) as: 1) Optics.Temperate_Early_Hardwood, 2) Optics.Temperate_Mid_Hardwood and 3) Optics.Temperate_Late_Hardwood. For each experiment, the simulations were run with multiple initial canopies to better examine the impacts of variation in the canopy structure.

Even though radiation availability affects production terms directly (e.g. photosynthetic carbon uptake), there are also longer-term impacts that occur, such as altering plant growth rates. This, in turn, is expected to change the canopy output terms over time as the vegetation structure changes as a consequence of the new light conditions. The variability and uncertainty in model outputs were thus evaluated at 1, 10, and, if necessary, 20 year increments, but only for the active growing season months (June 1st to October 31st) when deciduous trees typically have leaves in order to evaluate both the short and long-term canopy responses to variability in canopy RT parameterization.

Our model simulations and subsequent analyses focused on the impacts of uncertainties in the radiative parameters on model outputs. In this study we focused primarily on Net Primary Production (NPP) to represent vegetation growth, Albedo to represent surface radiation balance, and the Bowen ratio to represent the overall surface energy balance. In this study we selected NPP over aboveground biomass (AGB) as our focal response as we were more interested in evaluating the impacts of radiation parameter uncertainties on forest growth rate and dynamics than overall vegetation carbon stocks. We expected NPP to respond more strongly given its closer link to carbon uptake and respiration, which would lead to downstream changes in AGB as a result, and as such determined it was a better variable to explore the direct impacts of parameter uncertainties. Additionally there are generally multiple model processes and limitations determining biomass, while NPP is more directly driven by the radiation. We also provide results showing the impacts on the primary components of the carbon cycle in a more detailed case study: net ecosystem exchange (NEE) and Gross Primary Production (GPP), as well as autotrophic, heterotrophic and total <u>respiration.</u>

The uncertainty analysis was done by varying parameters at the quantile equivalents of −2, −1, 0, 1 and 2 Gaussian standard deviations (representing 2.3%, 15.9%, 50%, 84.1%, and 97.7% values). We modified the parameters across these quantile ranges using a univariate instead of multivariate change in parameter values for the sake of clarity. From these model simulations we then calculated model sensitivity, elasticity and the variance decomposition (LeBauer et al., 2013). Variance decompositions were then normalized by the median outputs to compare across different canopy types and simulations with different model output ranges.

Our first tests were done with synthetic canopies containing only one cohort at a time to isolate RTM parameter effects on individual PFTs and generate outputs similar to a so-called ‘big leaf’ model. In these tests, we assumed all trees were represented by a single cohort with an LAI of 6 and a diameter at breast height of 25 cm. The tests were done separately with cohorts of Early, Mid and Late Hardwood PFTs with stem densities of 0.0851, 0.0587 and 0.0682 stems per square meter, respectively. The stem density required to achieve constant LAI across the simulations varied among PFTs because of differences in SLA and foliar biomass allometries in the ED2 model (Medvigy et al., 2009; Medvigy et al., 2012).

Our next set of simulations were conducted with synthetic mixed canopies containing single Early, Mid and Late Hardwood cohorts with each contributing an LAI of 2, for a total LAI of 6. In this case the stem densities are 0.0284, 0.0196 and 0.0227 stems per square meter for Early, Mid and Late Hardwood PFTs, respectively. To further study the role of canopy structure in canopy radiative transfer modeling in ecosystem models, these simulations were done with two different canopies. The first canopy consists of three cohorts, all with 25 cm DBH and height order from tallest to shortest is Early, Mid and Late Hardwood (“EML”). In the second canopy the Early Hardwood cohort DBH was reduced to 20 cm to force the height order to be, from tallest to shortest, Mid, Late and Early Hardwood (“MLE”). For both canopies, the uncertainty analysis was done separately for each PFT to determine if the mixed canopy sensitivities can be approximated as a direct sum of individual PFT outputs or if the interactions within canopy are more complex.

Finally, the model uncertainty related to canopy radiative transfer was tested with simulations based on real forest inventory plots from the Willow Creek Ameriflux tower (45.8059 N, 90.0799 W) in Chequamegon-Nicolet National Forest in Northern Wisconsin, USA (Cook et al. 2004). This site is dominated by mature, evenly-aged deciduous hardwood forests. This test was conducted to examine the effect of increasing canopy complexity on the results and to evaluate the potential feedbacks from changes in canopy composition and structure due to changes in productivity, competition, mortality, and forest succession. For the sake of clarity and brevity, we only focus on the production terms and do not examine how the more complex canopy representation impacts our results of albedo and Bowen ratio. The inventory data were collected from a circular plot with a radius of 20 m and consisting of 2 Early Hardwood PFTs and 16 Late Hardwood PFTs. The inventoried cohorts are presented in Table 2. For the inventory canopy, we assumed that the stem density of each inventoried tree was one stem per area of the plot. We then analyzed the model sensitivity for the first, tenth and twentieth years of the simulation to test the impacts on model output at the beginning and throughout the simulation period.

**Table 2:** The forest inventory plot from Willow Creek, Wisconsin

| PFT | DBH [cm] |
| --- | --- |
| Late deciduous | 23.5 |
| Early deciduous | 5.3 |
| Late deciduous | 28.7 |
| Late deciduous | 8.6 |
| Late deciduous | 7.1 |
| Late deciduous | 26.4 |
| Late deciduous | 22.5 |
| Late deciduous | 8.9 |
| Late deciduous | 30.2 |
| Late deciduous | 30.4 |
| Late deciduous | 8.6 |
| Late deciduous | 20.4 |
| Late deciduous | 7.4 |
| Late deciduous | 29.6 |
| Early deciduous | 6 |
| Late deciduous | 28 |
| Late deciduous | 26.5 |
| Late deciduous | 23.7 |

To reduce the impact of the climate variability on the sensitivity results, we cycled the same climate drivers derived from 2004 meteorological data at the Willow Creek eddy covariance tower (US-WCr) through each year of the simulation (available from the Ameriflux data portal, http://ameriflux.lbl.gov/sites/siteinfo/US-WCr). The model simulations were conducted over twenty years with the first year starting from 1 June and the last year ending 31 December. The analyses include the output from both daytime and nighttime periods. However, it should be noted, that the nighttime shortwave radiation results are insensitive to the radiation parameters and properties given the lack of incident light. This may cause the variance and uncertainties to appear smaller, however given this was the default configuration for the uncertainty analyses in PEcAn we selected this approach for consistency with past work.

To simplify the comparisons between simulations across canopies, we normalized the results by the ED2 model output at the median parameter value. This is necessary as the model output will be different for a canopy consisting solely of Early or Late Hardwood PFTs even if their LAIs are the same. The normalization requires that the standard deviation is calculated from the variance decomposition. Finally, as the focus of this research was on how radiative uncertainties impact the model state instead of model validation, the resulting outputs will not be compared to measured values. Such an examination was not within the scope of the study here since the interest is rather on the radiative components affecting the total canopy output instead of what that total canopy output is.

## 3. Results

Below we present results highlighting the uncertainty in model projections of NPP, canopy albedo, and energy balance (Bowen ratio) with the ED2 model based on the prescribed variation of canopy RT parameters. In general, we observed a strong sensitivity of modeled fluxes and carbon stocks to RT parameters and resulting changes to canopy structure.

Figure 2 shows the normalized elasticity and the variance decomposition (VD, displayed as normalized standard deviation) for ED2 projections of NPP for each single cohort canopy at the first and tenth year of simulation. According to the VD results, the parameters driving uncertainties for a single cohort canopy, regardless of PFT, are clumping factor and leaf reflectance and transmittance in the PAR region (~400-700 nm). The elasticities also show that NPP increases with an increasing clumping factor (i.e. less clumped, more random leaf distribution) and decreases with larger leaf PAR transmittance and reflectance. For all three canopies the sensitivity for the RT parameters increases with the simulation time with Early and Mid Hardwood canopies displayed a higher sensitivity to RT parameters than the Late Hardwood canopy. As an example of how much these changes affect the total NPP of the canopy, changes in LT_PAR_ over the parameter range changed the total NPP for the simulated 10 years from ~ 285 MgC m^−2^yr^−1^ to 246 MgC m^−2^yr^−1^ (Decrease of ~14%) for the Early Hardwood canopy, from ~ 377 MgC m^−2^yr^−1^ to 322 MgC m^−2^yr^−1^ (Decrease of ~15%) for the Mid Hardwood canopy and from ~ 220 MgC m^−2^yr^−1^ to 208 MgC m^−2^yr^−1^ (Decrease of ~5%).

**Fig 2:**
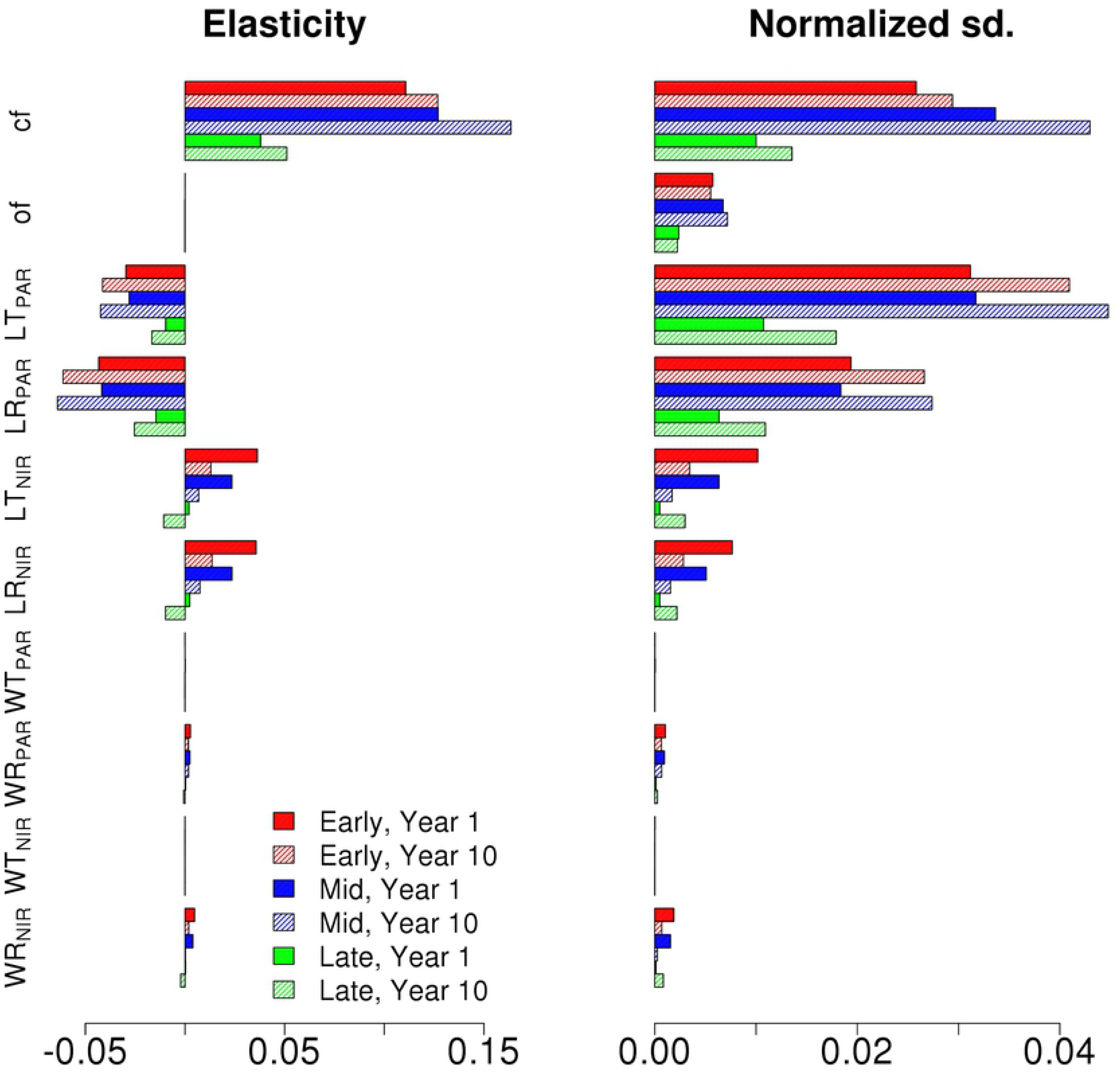
The NPP normalized elasticity and normalized output standard deviations of Early (Red), Mid (Blue) and Late (Green) Hardwood single cohort canopies for the chosen radiative parameters. Results are shown for first (full) and tenth (shaded) year of the simulation.

The elasticity and normalized VD for the single cohort canopy albedo are shown in Figure 3 for Early, Mid and Late Hardwoods compositions for both the first and tenth year of the simulation. The VD results highlight that although both PAR and NIR optical parameters affected albedo, parameters in the NIR domain have a larger impact on canopy albedo than the PAR range. The elasticities were positive for all parameters indicating that albedo increased with increasing parameter values (Figure 3). The albedo was initially more impacted by leaf parameter than wood, but over time albedo sensitivity to leaf parameters decreases while its sensitivity to wood parameters increases. Overall, we observed up to ~10% changes in albedo over our prescribed parameter uncertainty ranges (Figure 1).

**Fig 3:**
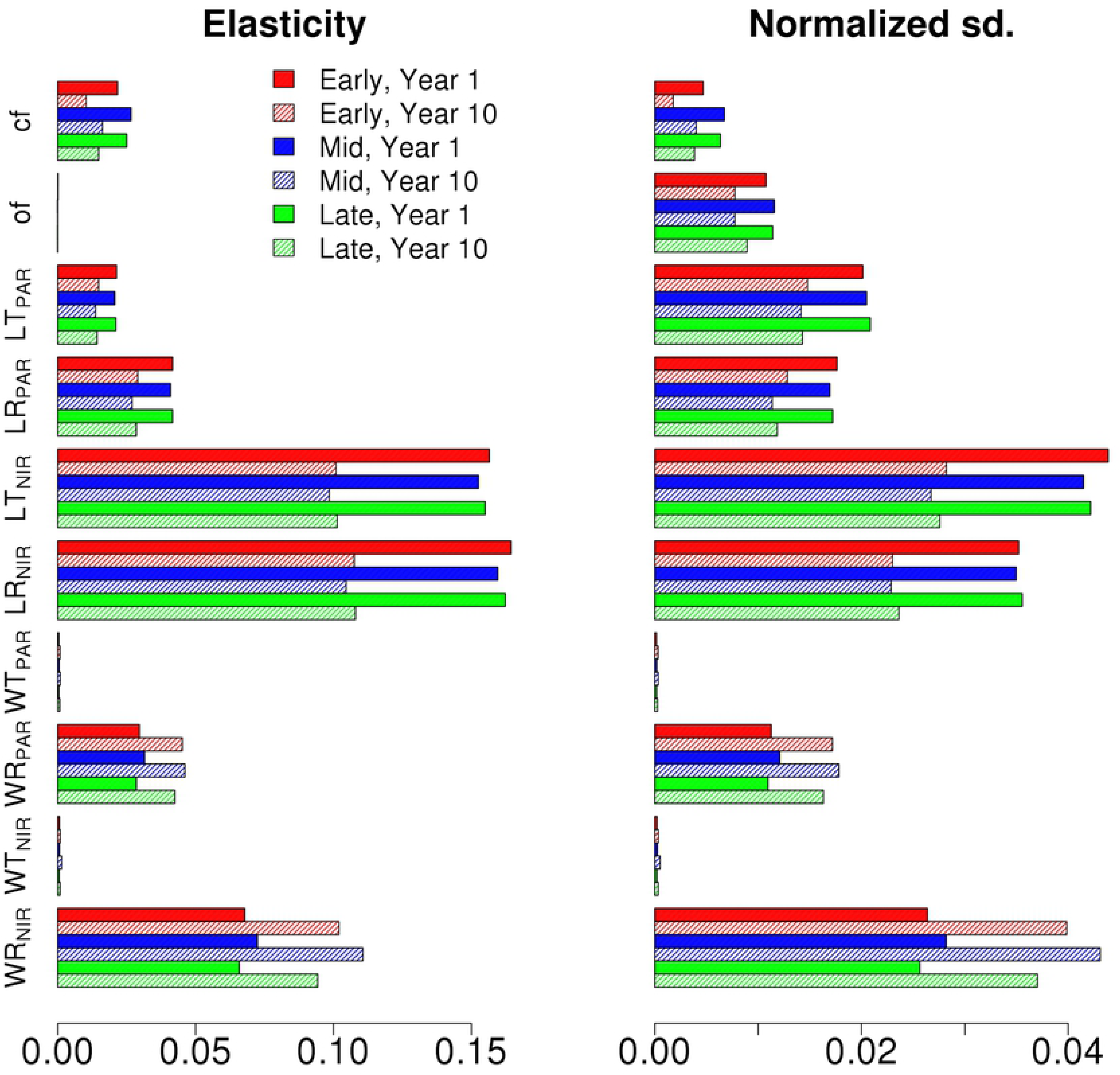
The Albedo elasticity and normalized output standard deviations of Early (Red), Mid (Blue) and Late (Green) Hardwood PFTs for the chosen radiative parameters. Results are shown for first (full) and tenth (shaded) year of the simulation

The elasticity and normalized standard deviation VD for the single cohort canopy Bowen show that the ED2 projected Bowen ratio for single cohort canopies is highly sensitive to wood reflectance (Supplemental Figure 1). In addition, the VD of both PAR and NIR wood reflectance parameters displayed an increasing trend over time, but importantly the direction of elasticity differed across the wood parameters indicating that increasing wood PAR/NIR reflectance decreases/increases the Bowen Ratio.

Figures 4 and 5 show the NPP elasticities and standard deviation decomposition for the different Hardwood PFTs for the synthetic mixed EML and MLE canopies over the first and the tenth years. From the VD results, it is evident that the uncertainties are driven by the tallest cohort in the canopy, in this case the Early Hardwood PFT in the EML (Figure 4) and Mid Hardwood PFT in MLE (Figure 5) canopies. In both cases, the clumping and orientation factors have the largest impact on production followed by LR_PAR_ and LT_PAR_. However, the uncertainties did not show a time dependence. It should be noted that when compared to the single cohort canopy elasticities in Fig. 2, the signs of PFT elasticities for the synthetic mixed canopies change based on their location within the canopy. In contrast to the single cohort canopies, NPP decreased with top cohort clumping factor and increased with top cohort leaf reflectance and transmittance. The NPP sensitivities for different hardwood PFTs in the synthetic mixed EML and MLE canopies for the first and tenth year of the simulation show that for clumping and orientation factors the responses to parameter values are non-linear. Additionally, for the EML stand, changing Early Hardwood leaf PAR transmittance over the parameter range increases the stand NPP from 279 MgCm^−2^yr^−1^ to 304 MgCm^−2^yr^−1^ (+19%) over the 10 year period while for the MLE stand changing the Mid Hardwood leaf PAR transmittance only increases stand NPP from 256 MgCm^−2^yr^−1^ to only 262 MgCm^−2^yr^−1^ (+2%).

**Fig 4:**
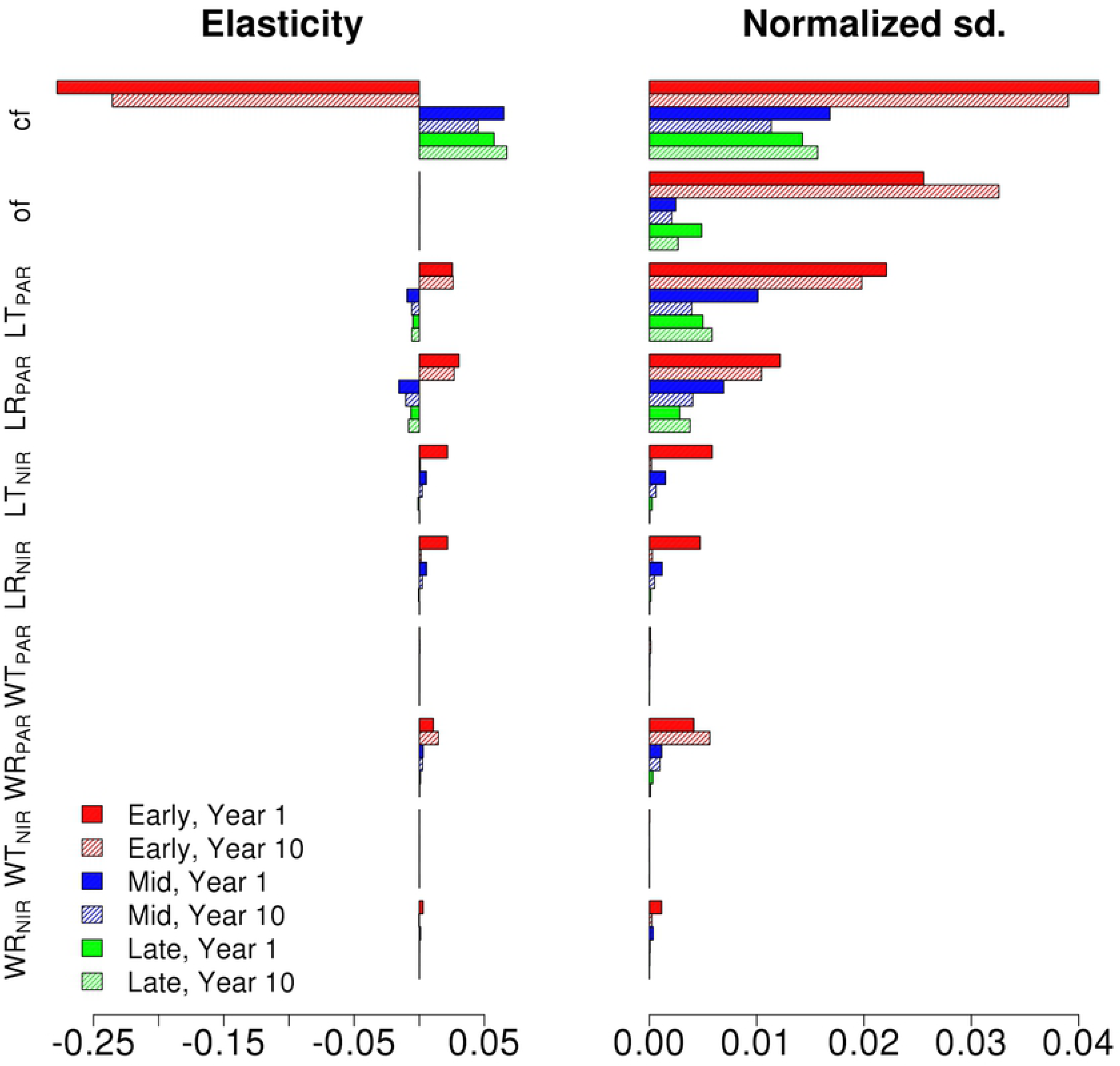
The NPP elasticity and normalized output standard deviations of Early (Red), Mid (Blue) and Late (Green) Hardwood PFTs in EML canopy for the chosen radiative parameters. Results are shown for first (full) and tenth (shaded) year of the simulation

**Fig 5:**
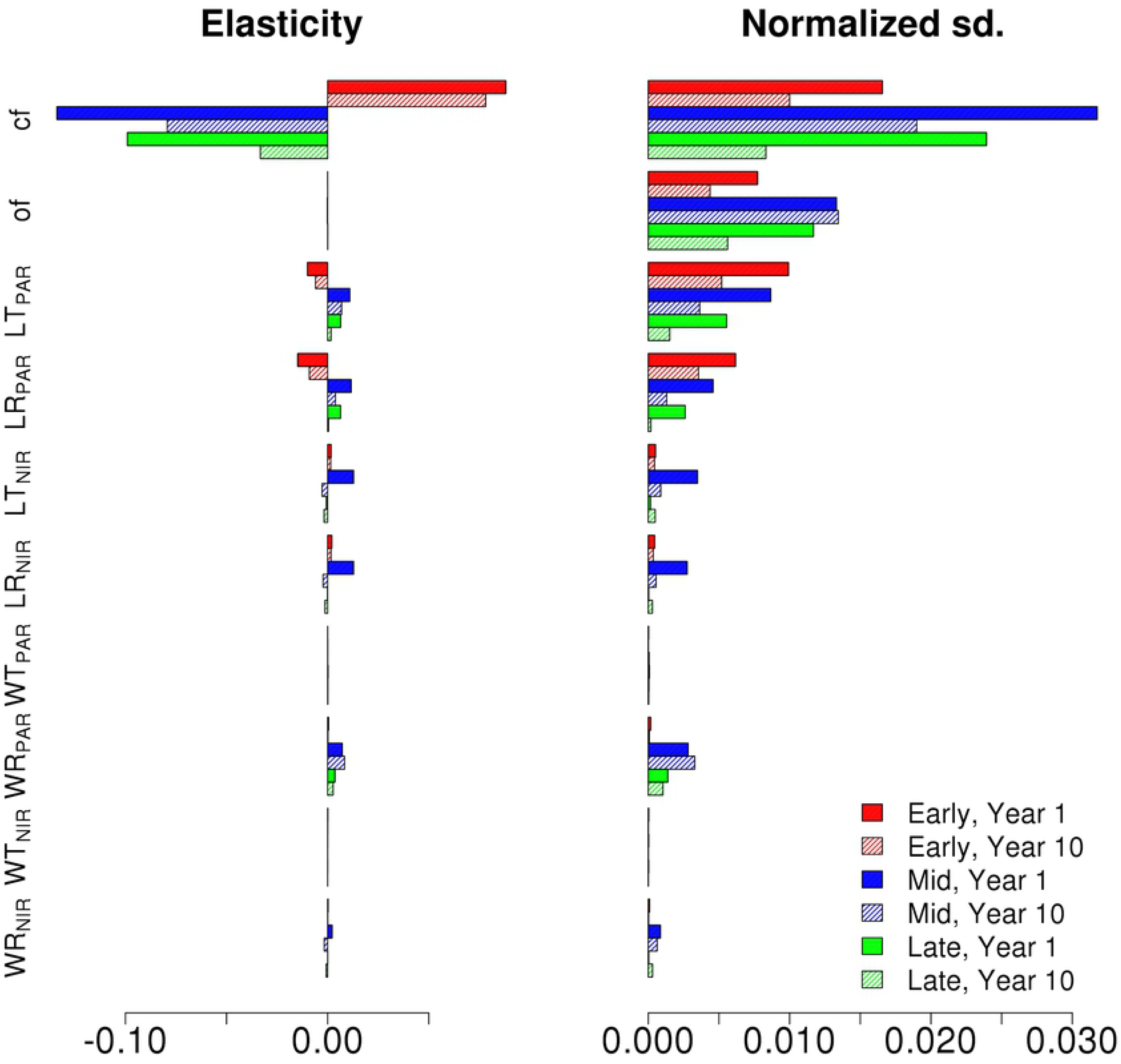
The NPP elasticity and normalized output standard deviations of Early (Red), Mid (Blue) and Late (Green) Hardwood PFTs in MLE canopy for the chosen radiative parameters. Results are shown for first (full) and tenth (shaded) year of the simulation

In Figures 6 and 7 we show the first and tenth year albedo elasticities and standard deviation decompositions for the different hardwood PFTs for EML and MLE canopies, respectively. The VD results show that the output uncertainty is driven by the parameter uncertainties in the tallest cohort. While the NIR parameters still drive the uncertainty, similar to Fig 3, the clumping and orientation factor affect the albedo more than for a single cohort canopy. The standard deviation decompositions decrease over time for the transmittance and reflectance parameters, but increase for the clumping and orientation factors. The albedo sensitivities for the synthetic mixed canopies for both the first and tenth year of the simulation show that while the albedo sensitivities vary based on PFT on the first year, the sensitivities are similar for all PFTs on the tenth year.

**Fig 6:**
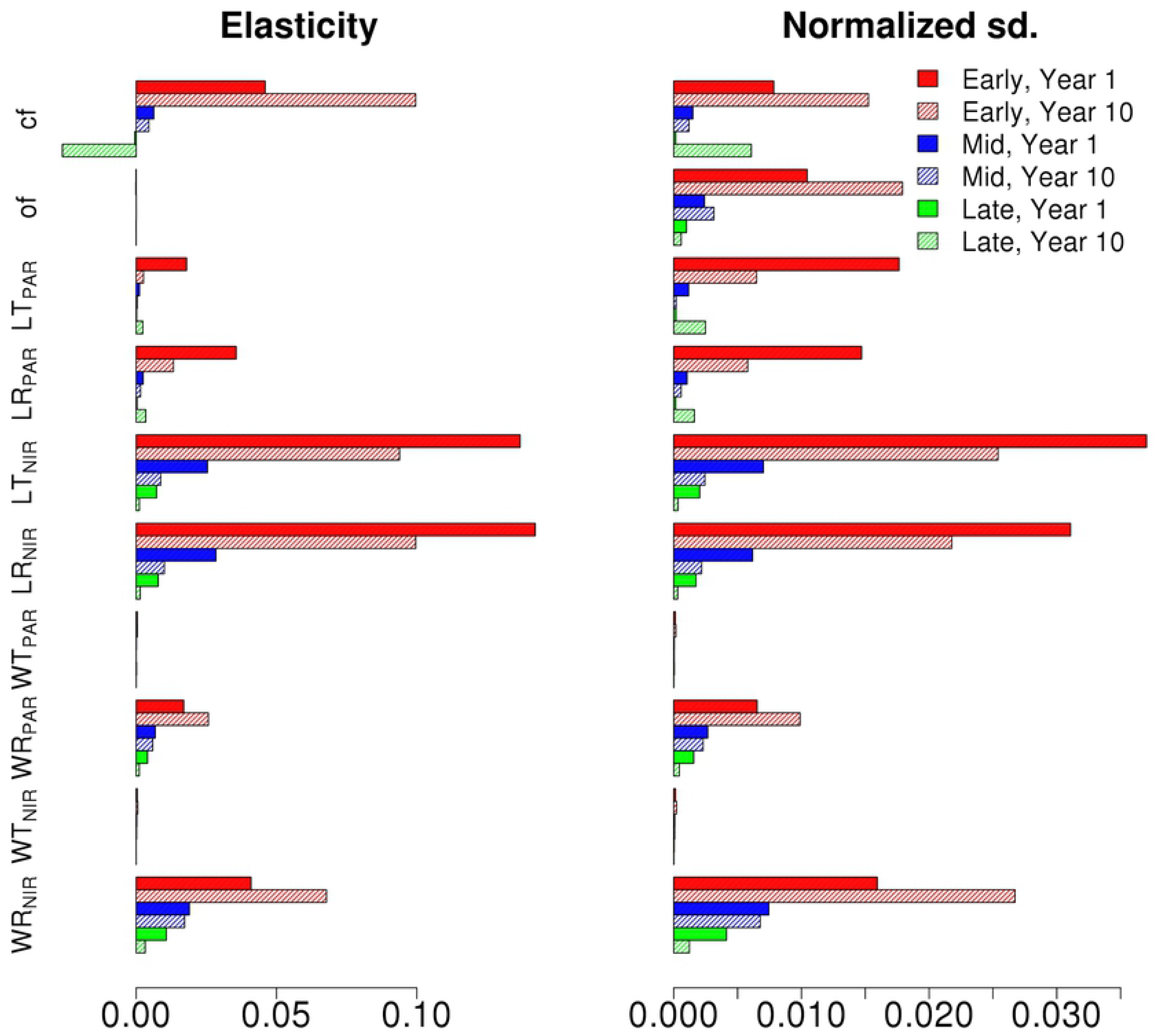
The Albedo elasticity and normalized output standard deviations of Early (Red), Mid (Blue) and Late (Green) Hardwood PFTs in EML canopy for the chosen radiative parameters. Results are shown for first (full) and tenth (shaded) year of the simulation

**Fig 7:**
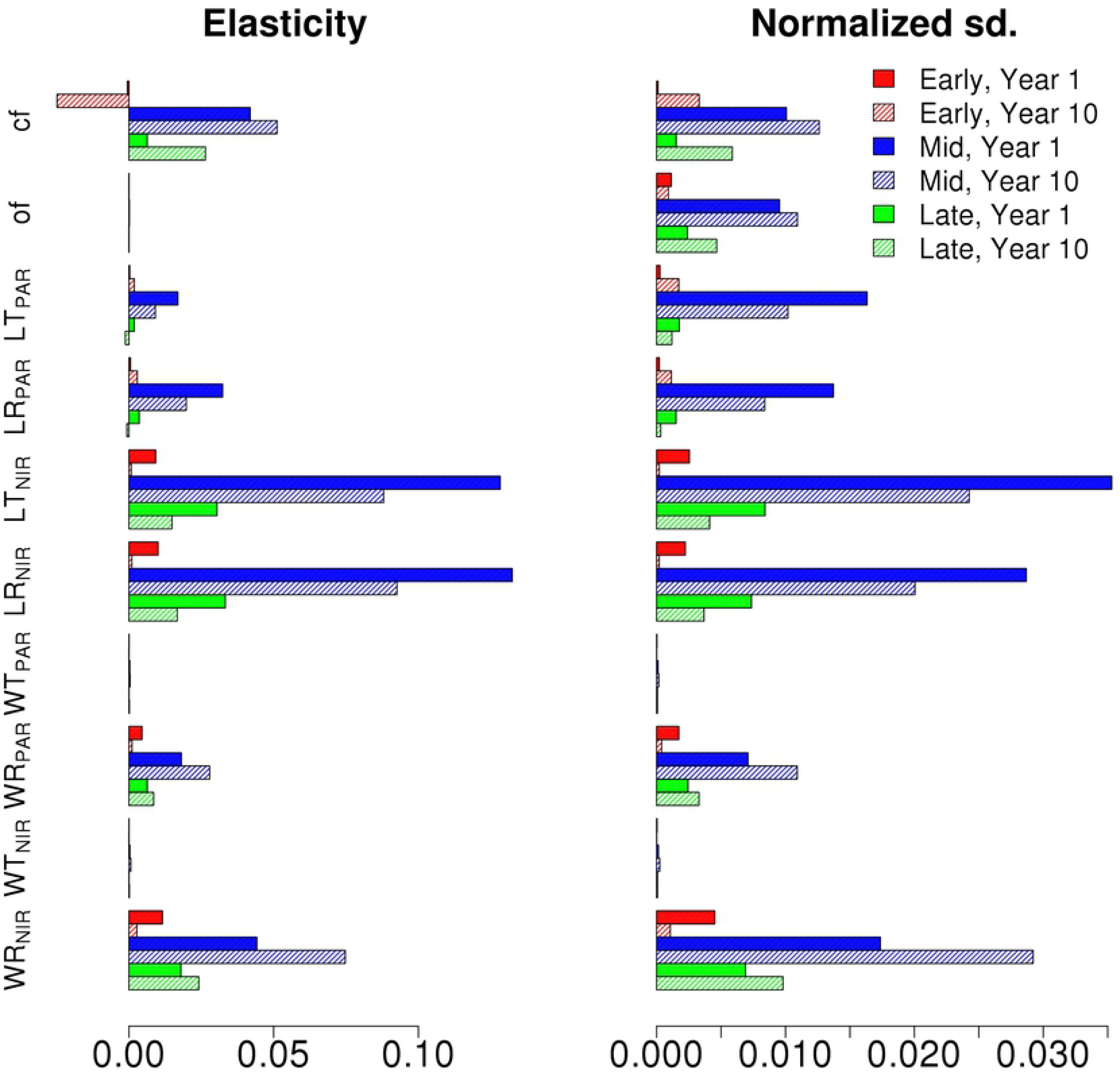
The Albedo elasticity and normalized output standard deviations of Early (Red), Mid (Blue) and Late (Green) Hardwood PFTs in MLE canopy for the chosen radiative parameters. Results are shown for first (full) and tenth (shaded) year of the simulation

The Bowen ratio elasticities and standard deviation decompositions for the different Hardwood PFTs are shown for the synthetic mixed EML and MLE canopies for the first and the tenth years are presented in Figures 8 and 9, respectively. The Bowen ratio of the canopy is shown to be dominated by the tallest cohort in the canopy and, similarly to single cohort canopies, the wood reflective properties have a notable effect on the Bowen ratio which increases over time. However, here the leaf radiative properties show large normalized deviations for the first year of the simulation, but decrease drastically over the simulation.

**Fig 8:**
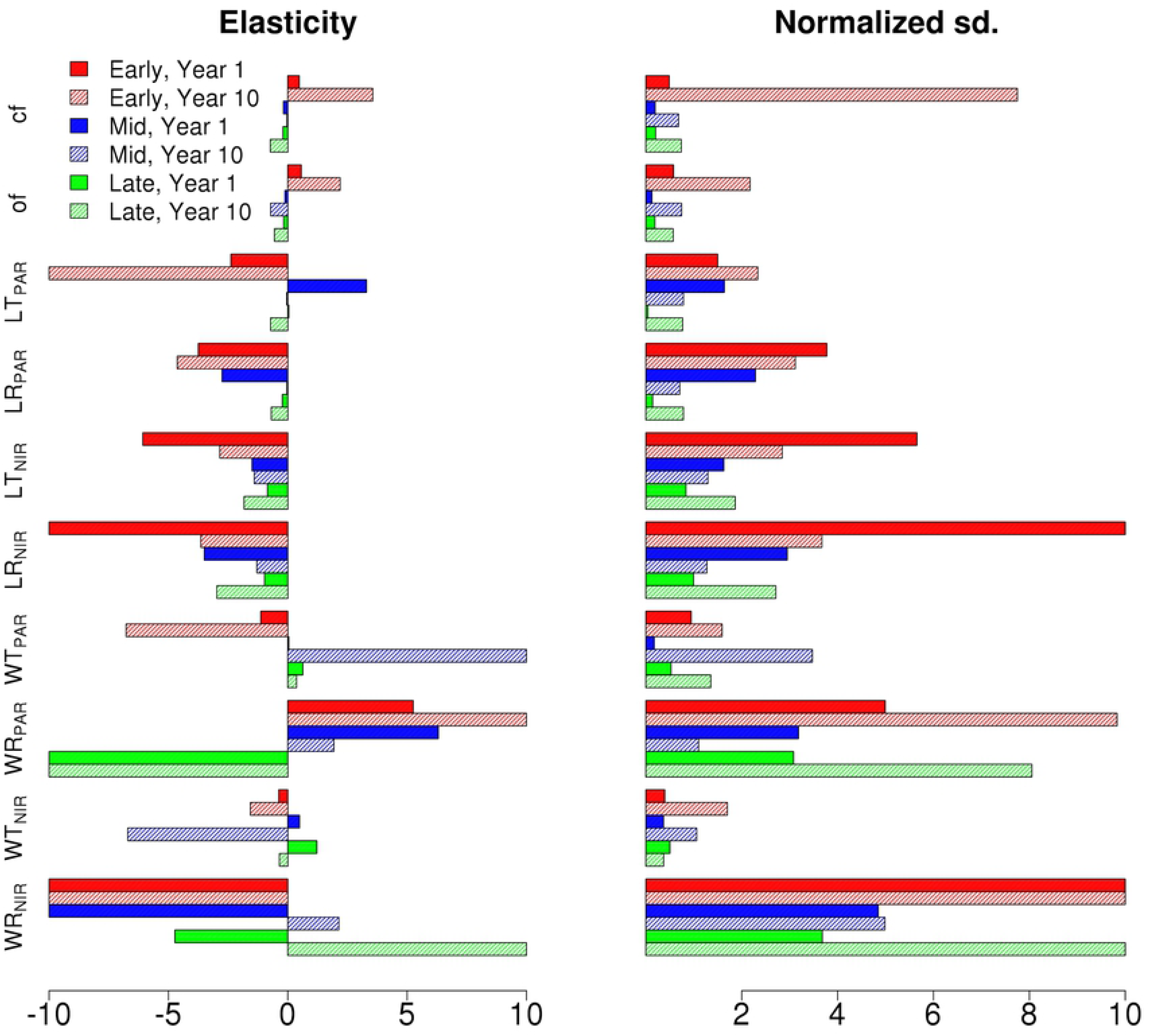
The Bowen Ratio elasticity and normalized output standard deviations of Early (Red), Mid (Blue) and Late (Green) Hardwood PFTs in EML canopy for the chosen radiative parameters. Results are shown for first (full) and tenth (shaded) year of the simulation. The elasticities were capped of at 10 in order to better compare the decompositions.

**Fig 9:**
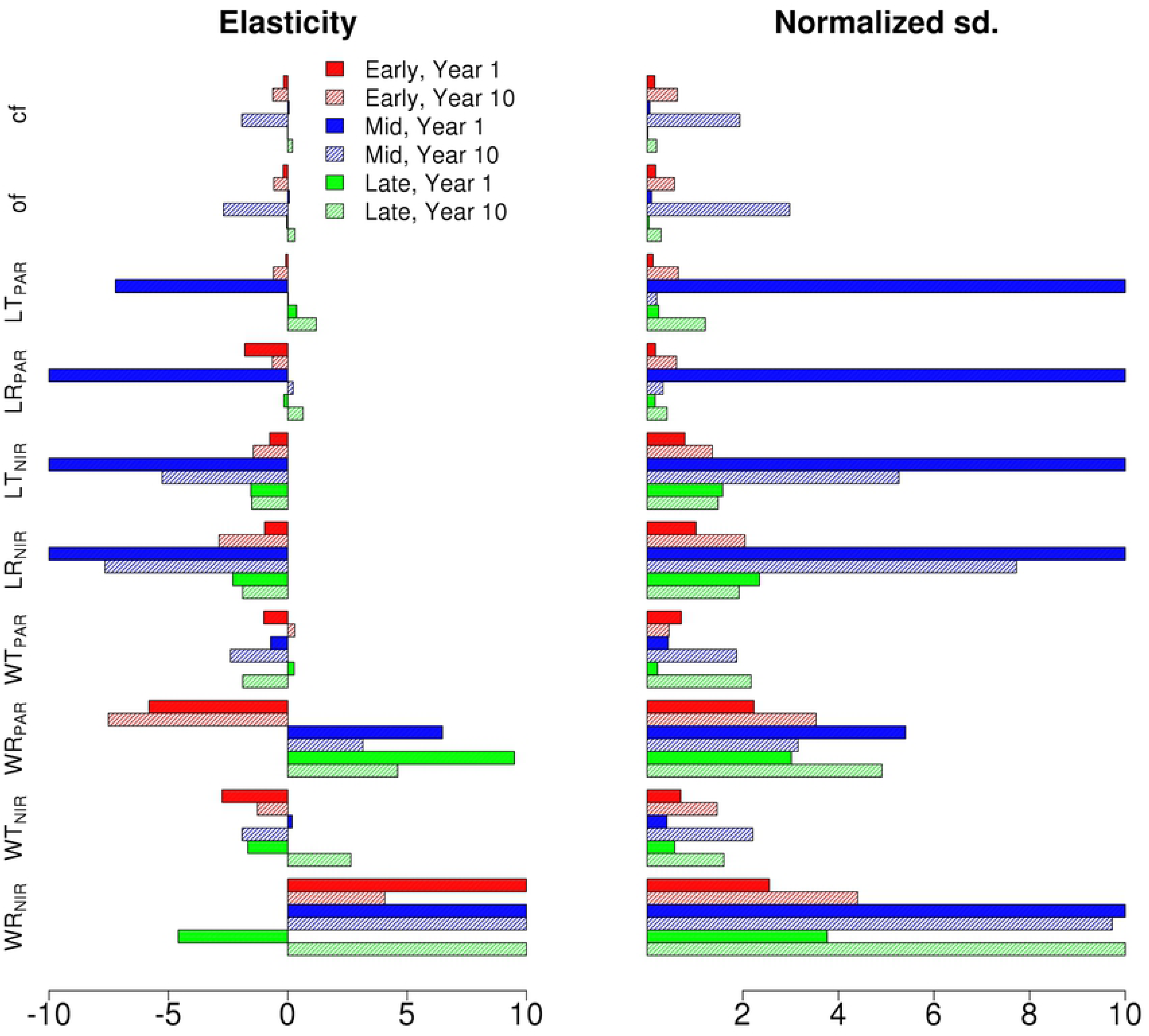
The Bowen Ratio elasticity and normalized output standard deviations of Early (Red), Mid (Blue) and Late (Green) Hardwood PFTs in MLE canopy for the chosen radiative parameters. Results are shown for first (full) and tenth (shaded) year of the simulation. The elasticities were capped of at 10 in order to better compare the decompositions.

To assess how changes in canopy radiation parameter values affect the resulting canopy structure through time, especially understory growth and regeneration, we present Fig.10, which shows how the proportion of biomass attributed to the tallest cohort in year 10 varies as a function of LT_PAR_ and clumping factor in the EML and MLE canopies. Changes in Early Hardwood parameter values in the EML canopy causes the largest changes in composition, with top cohort fractional abundance decreasing by almost 25 percent with increasing LT_PAR_. The MLE canopy changes in structure are more subtle, with the proportional amount of biomass in the understory increasing by 5%. For both canopies, increases in the understory biomass correspond with an increase in total canopy biomass while the tallest cohort biomass remains approximately the same (not shown). We note that the differences in biomass distribution increase with longer simulations (results from the first year not shown).

**Fig 10:**
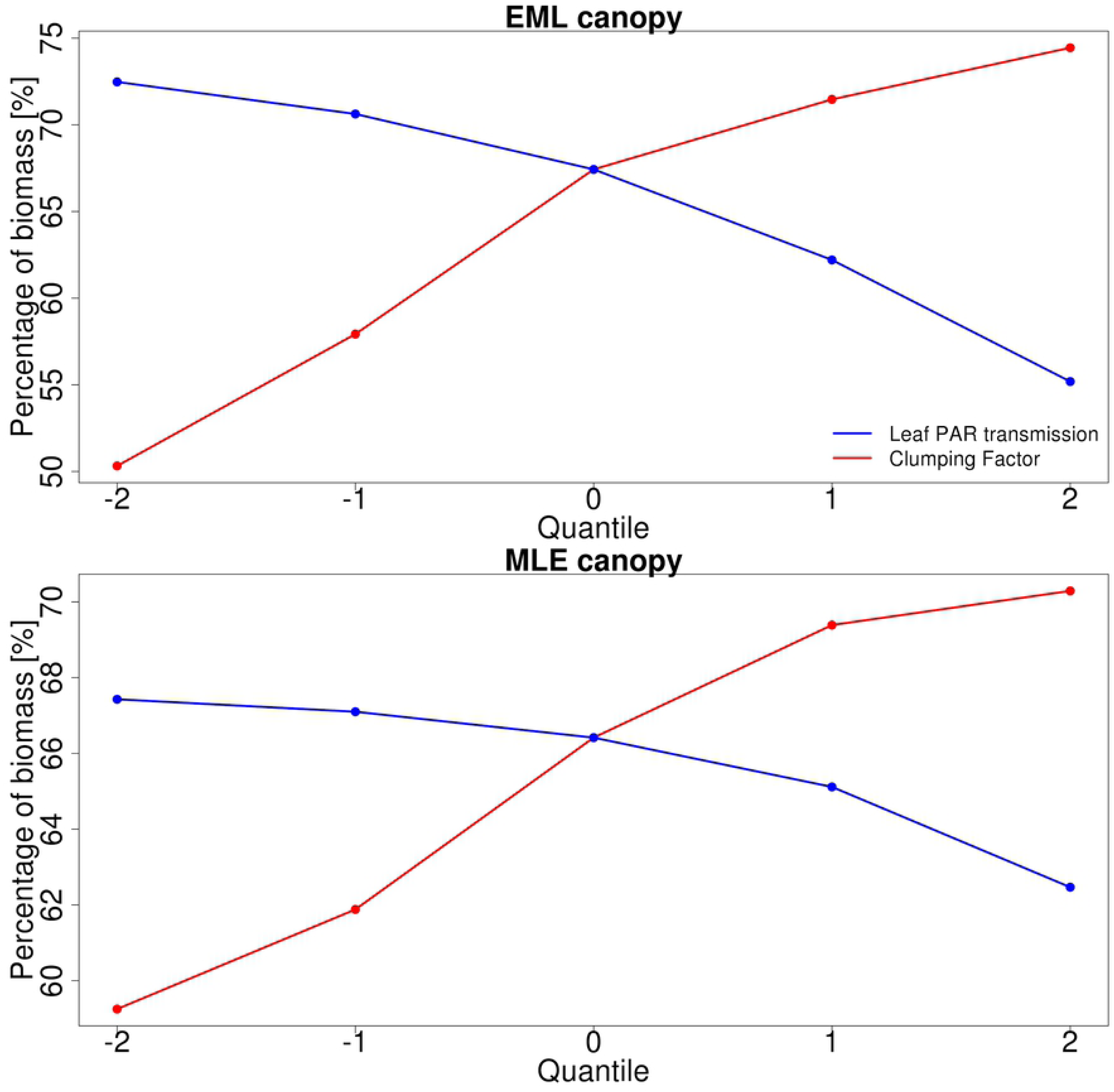
The sensitivity of the percentage of tallest cohort biomass from total canopy biomass to both leaf PAR transmission and clumping factor. The upper figure is for the EML canopy and the sensitivities are for Early Hardwood PFT and the lower figure is for the MLE canopy and the sensitivities are for Mid Hardwood PFT.

When utilizing Willow Creek inventory data as model initial conditions, we find again that clumping and orientation factor dominate uncertainty (Supplemental Figure 2) for the NPP of the Late Hardwood PFT NPP. Moreover we found that variation in leaf PAR transmittance and reflectance had a smaller impact on the total canopy NPP than with the single or mixed cohort canopies (Figs. 2, 4 and 5). The variable decompositions of the model outputs in response to changes in Late Hardwood LT_PAR_ (Supplemental Figure 3) show that NEE was most affected by changes in parameter values, but the resulting standard deviation decreases over time as respiration and carbon uptake and storage (GPP and NPP) increase.

## 4. Discussion

Our results establish that radiative parameter uncertainties can have considerable impacts when projecting carbon, water and energy fluxes with TBMs that deploy the two-stream approach and which contain a demographic representation of vegetation dynamics. This finding is in agreement with previous studies evaluating the influence of canopy radiation transfer (RT) in other TBMs (e.g. Alton et al., 2007; Koybayashi et al., 2012). Moreover, we highlight how canopy RT uncertainties affect canopy carbon allocation, which in turn would influence terrestrial carbon cycle projections (Friedlingstein et al., 2014). As the two-stream approach is used in a number of widely-used models that also represent the land component of Earth System Models (ESMs), including JULES (Best et al., 2011) and CLM (Lawrence and Fisher, 2013), these results underline the importance of properly constraining the parameters that regulate the canopy radiation regime across different vegetation types and biomes, as has been demonstrated for other associated processes (e.g. Dietze et al., 2014; Rogers et al., 2017). In addition, related processes such as succession, competition, recruitment, and mortality in a demographic model, such as ED2, may similarly be directly and indirectly impacted by canopy RT uncertainty (Fisher et al 2017). The direct impacts result from changes in the stand-level carbon balance driven by light utilization and surface temperature and indirectly from changes in water balance as well as longer term changes in internal composition and structure. These competitive interactions complicate the interpretation of the canopy response to radiation parameter values, particularly within models such as ED2 that represent the canopy as a series of cohorts across a size and age class distribution (Fisher et al 2017), as we observed in this study. As a result more careful evaluation of the combined uncertainties associated with canopy RT, growth, mortality, allometry and competition is needed but was beyond the scope of this work.

Our evaluation of TBM projection uncertainty is relevant to the larger TBM and ESM communities for several key reasons. Many TBMs utilize a variant of the two-stream model (Sellers et al., 1992) to calculate canopy radiative fluxes and the leaf / canopy light harvesting to drive photosynthesis, energy balance, and associated processes such as transpiration and decomposition. All of these interactions, however, need to be considered during model development as examining individual model outputs that represent the aggregate stand-level response, such as the total canopy GPP or NPP (as opposed to the cohort or PFT-scale values), could result in misleading conclusions given the complex internal model dynamics.

It has already been established that the choice of RTM and the parameterization affects the production outputs of TBMs, with both big-leaf and detailed 3-D canopy architectures (e.g. Alton; 2007; Bonan et al., 2011; Chen 2012; Kobayashi et al., 2012). Our results expand on this by showing that not only do the uncertainties in radiation parameters impact demographic TBMs such as ED2 projections, but also that these impacts depend strongly on canopy structure. Our results show that the uncertainty in model projections of NPP (Figs 2, 4 and 5) is large (up to 20 %;), even over the relatively short simulation period here, with respect to realistic variation in canopy RT parameterization for a temperate broadleaf deciduous forest ecosystem. This uncertainty also showed strong variation through time as a result of changes in canopy structure, internal model dynamics, and competition/succession of cohorts; including complex vertical changes resulting in a canopy structure that was in stark contrast to reality, as observed in the US-WCr site inventory data, for example relating to the mortality/survival of stems in the understory. In the simulations initialized with the inventory data, the shortest cohorts were so light-starved that their stem density constantly decreased over time indicating cohort mortality in situations were in reality they survived and grew.

The complex impact of the structure on the radiation parameter sensitivity, and consequently the possibility for misleading conclusions, can be seen when comparing how the NPP parameter sensitivity develops over time for single cohort (Fig. 2) and multi-cohort (Figs. 5 and 6) canopies. In the single cohort canopies all the stems are of the same height, so there is no competition for radiation and all of the cohorts have the same NPP. Here if the cohorts absorb more radiation, they also have increased growth, and vice versa. As the output sensitivity at a given time is determined by comparing the modeled canopy outputs at that time and the larger cohorts are also able to absorb more radiation, this results in the canopy NPP sensitivity for the central radiation parameter values becoming more pronounced over time. The single PFT canopy results also highlight how important cohort PFTs are not only for determining how sensitive the cohort is to radiative parameter uncertainties, but also on how they respond to changes in radiation absorption. The Late Hardwood NPP sensitivity for LT_PAR_ (Fig 2) is a good example of this as while it is least sensitive of the tree PFTs to LT_PAR_, the sensitivity also has the largest relative increase over time.

In the more complex EML and MLE canopies, though, the NPP sensitivities appear to decrease over time (Figs 5 and 6). When the radiation absorption of the tallest cohort is reduced, the decrease of that cohort’s NPP will cause the total canopy NPP also to decrease. However this allows there to be more radiation available in the understory which in turn will enhance growth and cohort survival there. As a result the understory NPP increases over time, partially balancing out the negative change in total canopy NPP. Similarly increasing the radiation absorption of the tallest cohort will increase its NPP, but the reduced radiation availability in the understory, and the consequent reduced growth and cohort survival, will lessen the positive total canopy NPP change over time. Consequently when examining the canopy NPP over time, the canopy can appear to be insensitive to radiative parameter while in actuality the results reflect the canopy structure adapting to the new radiation availability.

Recent efforts have focused on the inclusion of layered canopies and cohort demographics into other ecosystem models (e.g. CLM(ED); Fisher et al., 2015; now FATES, https://github.com/NGEET/fates). However, based on our results, the radiative parameter values play a crucial role in understory growth and cohort survival as they dictate the amount of available radiation for the understory cohorts. This supports previous studies that show the importance of capturing canopy structure to improve model predictions (e.g. Antonarakis et al., 2011), but also highlights how sensitive the model results are to canopy structure or changes in structure through time. Therefore, it is important to not only accurately identify the forest composition or plant functional types (PFTs) of the trees within the canopy, but also how they interact within the canopy layers (i.e. competition for light), as represented by the radiative parameters, particularly with demographic models such as ED2 (Fisher et al., 2017). Importantly for the basic ED2 internal canopy radiative transfer model the canopy albedo is dominated by the uncertainties of the tallest cohort, indicating that increasing the number of cohorts or more accurately representing understory structure will not necessarily improve the projections and evolution of canopy albedo.

When examining changes in total canopy biomass (not shown), the response to radiative transfer model parameter value changes were considerably weaker, or even absent with the more complex canopies. This was in sharp contrast to NPP but not unexpected as biomass represents the overall carbon pools while NPP represents the change in biomass or growth through time, and is much more closely linked with photosynthesis over the short term than Above Ground Biomass (AGB); the amount of standing live and dead biomass is regulated by a number of factors including the rates of carbon uptake and losses, as well as turnover processes. Additionally AGB is more constrained by internal model assumptions as, for example, ecosystem models limit how much a cohort can grow in a year to avoid unlimited growth. However in the tests here the tallest cohorts are so light-saturated that while their NPP will change in response to available radiation, it will still be so high that the top cohort AGB will only incrementally change.

The impact of the radiative parameter values on the biomass distribution within the canopy (Fig 10) highlights both the factors discussed before while also further illustrating the role the radiation parameters play in how the canopy structure develops over time. While the canopy NPP is most sensitive to the radiative properties of the top cohort, the changes in total canopy AGB are almost completely driven by the understory AGB. This is due to the light-limited understory which causes the biomass there to be competitive over the remaining radiation and also causes it to react more strongly to changes in radiation availability (Bond-Lamberty et al., 2015). Not only do we see how the understory growth/diminishing depends on canopy structure, as indicated by the differences between the biomass responses of the EML and MLE canopies, the survival of the shortest cohorts, which are most starving for radiation, is strongly connected to the canopy radiation parameter values (not shown). Additionally these results show that the accumulation of biomass in the understory in response to changing radiation conditions strongly depends on the PFT composition there.

The parameterization uncertainty of the two-stream model in ED2 also lead to significant variability in the projections of the energy cycle, where albedo (Figs 3, 6 and 7) and the Bowen ratio (Figs S1, 8 and 9) showed strong variability in model output. Alton et al. (2007) found a similar response with the JULES model, and together these results show the importance of improving the representation of canopy RT parameterization and structure in TBMs as the carbon, water and energy cycles all play a key role in regulating climate (Bonan, 2008). Our results and analyses highlight several important parameters to consider and constrain when exploring ways to improve TBM fidelity. In particular surface albedo is a key driver in the surface to atmosphere energy balance, which is why it is important to correctly determine how both optical parameter uncertainties and canopy structure influence the modeled canopy albedo (Kaminski et al., 2016). However, it is also important to consider how albedo is defined. Here, albedo was considered as the total surface albedo covering the combined PAR and NIR radiation range (400 to 2500 nm). Plants reflect the most (while absorbing the least) radiation in the NIR, followed by the two peaks in the shortwave infrared (Ollinger, 2011). Consequently the parameters controlling the NIR reflectivity dominated the ED2 albedo, especially the wood NIR properties as the leaf NIR parameter values are so small and vary so little that they do not really have impact on the albedo (Fig 1). What is noteworthy, however, is that while clumping factor and leaf PAR transmission or reflectance can have a strong effect on growth, in our analysis their impact on albedo was more muted.

Thus optimizing the model for one response, such as the total canopy albedo, will not necessarily improve the representation of other processes in the model, for example how well it simulates carbon uptake and storage. As a result, when optimizing model parameters during model development, it should be conducted with multiple datasets for multiple responses rather than just relying on a single information source. Also, while the sensitivities remain relatively constant over time with single cohort simulations, there are individual PFT sensitivities that clearly show different responses that also evolve through time (See Fig.7). These results underline how important it is to consider the PFT composition of the canopy when modeling radiation.

The thermal energy regime of vegetation canopies can impact the surface energy and fluxes on the local, regional, and global climate scales (Amiro et al., 2001; Bonan, 2008). In our study we found strong variation in the modeled Bowen ratios in response to changes in canopy RT parameters, especially with respect to the WR_PAR_ and WR_NIR_. The Bowen ratios produced by the median parameter values are within reasonable ranges for a deciduous forests (Wilson and Baldocchi, 2000), but evolved rapidly (Fig 8) over reasonable input parameter values (Fig 1). The impact of leaf parameters on the mixed canopy Bowen Ratio lessened over time, implying that the canopy adjusted to the new state, while the wood parameter impacts increased over time, indicating the model did not balance out changes in wood reflectance parameter values during the length of our simulations. These effects are possibly due to the representation of the vertical canopy temperature profile in the model (or lack thereof) as previous studies have shown that that leads to unrealistic sensible heat fluxes (Baldocchi and Wilson, 2001).

These internal dynamics can easily make mixed canopies appear insensitive to parameter variation. For example, the single cohort simulations showed that leaf PAR transmittance and reflectance had a straight-forward impact on the canopy NPP, however with the synthetic mixed canopies those responses become more complex (Figures 2, 4 and 5). Furthermore, our simulations using measured plot inventory initial conditions showed that, on the whole, canopy NPP did not show a strong response to parameter uncertainties. However, further analysis showed that the individual cohort growth within the canopy varied strongly in response to leaf PAR transmittance and reflectance, while respiration and NEE showed consistent changes in response to changes in leaf PAR transmittance (Figures 3, 6 and 7). Therefore if the majority of the canopy consists of the same PFT, as it had with our inventoried canopy, it can result in a total stand-level NPP that is fairly static but the internal distribution of NPP across cohorts can be much more dynamic. This further highlights how using just the total canopy outputs to examine and to constrain the canopy radiation parameters in demographic models can very easily lead to misleading results in height-structured models.

While the focus of this paper was on the radiative transfer parameters, the results also strongly highlight the interactions between the canopy state and radiation profile. The allometric relationships determining the foliar biomasses, and thus LAI, have their own associated uncertainties which also affect the output variables. In addition, the radiation distribution within the canopy plays an important role in the growth and mortality of cohorts. All of this indicates that the effect of allometric uncertainties should be studied more and included in the projection uncertainties.

From the perspective of TBMs, it is important to emphasize that radiative processes are a central part of all TBMs as they govern how much radiation is available for the vegetation and, consequently, are a key model driver. As other processes have been included and calibrated, their outputs have been impacted by the assumptions made about the available radiation in different parts of the canopy. Thus, when examining the radiative parameters, we found that PFT variation can be large in response to realistic radiation values. The radiative parameters cannot be calibrated simply against measurements without also taking into account how the models themselves are calibrated based on those radiative properties. Thus if other processes have been calibrated with erroneous radiation parameters or process representation, more strongly constraining the radiative parameter values could potentially reduce the model accuracy if those other parameters were not recalibrated accordingly.

Looking forward, as radiation is a central driver for the ecosystem models, tightly constraining radiative parameters in TBMs should be a priority due to their associated uncertainties affecting key outputs relating to growth and energy balance as shown by the results here. Furthermore the canopy structure is a crucial component in canopy response and adjustment to changing radiation availability, the canopy structure representation and the radiation scheme will always be linked and thus need to be developed together. Finally different RTMs should also be tested according to the framework presented here in order to determine how much their associated radiation parameter uncertainties impact the model state.

### Limitations of the present study

We did not consider the covariance in parameter values, and instead evaluated model responses to single parameter changes at time to simplify the analysis. However, future work should also consider the complex interactions between parameters on the resulting model behavior, but this was beyond the scope of this study.

Additionally we focused on model outputs over the entire day (i.e. full light-dark cycle). However, without solar radiation, the canopy outputs are insensitive to the SW radiation parameters, which likely leads to an underestimate of the model elasticity.

## 5. Conclusions

Our results show that radiative parameter uncertainties impact the model output results for canopy production, radiation balance, internal energy, and demography. Our work not only establishes the importance of constraining optical parameter values with data sets such as remote sensing at the leaf to landscape scales, but also that constraining the model with, for example, albedo observations will not necessarily improve predictions on productivity. Instead multiple data constraints at different scales are needed to inform model projections as well as to test different model representations of canopy radiation transfer and demography. This aspect needs to be accounted for when assessing model performance.

Additionally we found a strong dependence on canopy structure. The change in internal radiation profile causes individual cohorts to have offsetting responses, such as increase NPP for some cohorts while decreasing it to others, which causes complex canopies appear to be insensitive to changes in radiation parameters even if they develop in different manners. This could impact how canopies grow in models such as ED2 and result in incorrect canopy succession and structure which in turn could lead to erroneous estimates of the C, water, and energy balances and fluxes. The influence of canopy structure as highlighted here shows that the importance of accurately simulating the initial canopy, but also the need to further develop radiation modeling, especially concerning how layered canopies are structured and implemented within TBMs. As new TBMs containing a demographic representation of canopy dynamics become available the issues raised here will become more important and need further attention to minimize potential impacts on modeling efforts.

## Acknowledgements

This study was funded by the National Aeronautics and Space Administration (NASA) through the grant NNX14AH65G. Additional support was provided by the United States Department of Energy contract No. DE-SC00112704 to Brookhaven National Laboratory and by NASA Grant 16-EARTH16F-0126 and NSF 1458021 to Boston University.

The model settings and simulation outputs used here are available from https://github.com/TESTgroup-BNL/Viskari_etal_ED2_radiation_uncertainty.

